# Quantitative Analysis of Trans-synaptic Protein Alignment

**DOI:** 10.1101/573063

**Authors:** Jia-Hui Chen, Thomas A Blanpied, Ai-Hui Tang

## Abstract

Nanoscale distribution of proteins and their relative positioning within a defined subcellular region are key to their physiological functions. Thanks to the super-resolution imaging methods, especially the single-molecule localization microscopy (SMLM), mapping the three-dimensional distribution of multiple proteins has been easier and more efficient than ever. In spite of the many tools available for efficient localization detection and image rendering, it has been a challenge to quantitatively analyze the 3D distribution and relative positioning of proteins in these SMLM data. Here, using the heterogeneously distributed synaptic proteins as examples, we describe in detail a series of analytical methods including detection of nanoscale density clusters, quantification of the trans-synaptic alignment between these protein densities, and automatic enface projection and averaging. These analyses were performed within customized Matlab routines and we make the full scripts available. The concepts behind these analytical methods and the scripts can be adapted for quantitative analysis of spatial organization of other macromolecular complexes.

**Highlights:**

- **Localization microscopy provides sufficient data for precise quantitative analysis.**
- **An algorithm to identify local density peaks within a 3D localization cluster.**
- **New methods for quantitative analysis of trans-synaptic proteins alignment and enrichment.**
- **These algorithms can be easily adapted to analysis of other subcellular organizations.**

## 1. Introduction

Neuronal communication via synaptic transmission is a complex biological process that must coordinate specialized protein structures within connected cells. Proteins in the presynaptic active zone mediate the release of neurotransmitters which diffuse within the synaptic cleft and activate postsynaptic receptors[1]. In spite of their significant impacts on synaptic transmission[2–5], subsynaptic structure and protein interactions are still unclear. This is chiefly because synapses are too small (hundreds of nanometers in diameter), existing beyond the limitation of optical diffraction of the conventional light microscope. Electron microscopy provides high enough resolution[6], but it is difficult to specifically mark and accurately recognize particular protein species, and cannot be applied in living cells. Recently developed methods of single molecule localization microscopy (SMLM) [7–9] provides the best opportunity for visualizing the protein distribution at structures such as synapses.

SMLM includes a variety of super-resolution imaging techniques that localize isolated fluorescent molecules with precision well beyond the diffraction limit by fitting their images with a version of the microscope’s point spread function. Application of these methods, especially stochastic optical reconstruction microscopy (STORM)[9,10], photoactivated localization microscopy (PALM)[8,11] and point accumulation for imaging in nanoscale topography (PAINT)[12], has led to series of discoveries of new biological structures and processes[13–18]. Specifically in neuroscience, these methods have revealed a new layer of protein organization at nanometer scale that are critical for modulation of synaptic functions. Postsynaptic scaffolding proteins are organized in nanoclusters enriched with AMPA receptors[19,20], while the presynaptic vesicle fusion sites are guided by nanoclusters of active zone proteins RIM [21] and Munc13[22]. Most surprisingly, nanoclusters of postsynaptic scaffolds and receptors were found to spatially aligned with presynaptic RIM nanoclusters, suggesting a trans-synaptic nanocolumn structure[21] that couples presynaptic transmitter release to the densities of postsynaptic receptors and optimizes the synaptic transmission[4,5,19,23]. The reorganization of nanocolumns in synapses may underlie the tuning of synaptic strength during plasticity and pathological conditions[4,24].

Despite the magnificent details SMLM has provided, performing quantitative analysis on this data has proven to be a challenge. This has become a barrier for the efficient application of SMLM in the biomedical field, especially since more sophisticated and customized analyses are often required to meet the demands of most specific projects than the image processing capabilities of most general software packages[25]. Indeed, many reports rely heavily simply on presenting images rather than exploiting the wealth of information present in them. Here, we describe detailed analytical methods on quantification of trans-synaptic alignment on three-dimensional STORM data with the full Matlab script attached. Some of the methods could be easily adapted to analysis of other biological structures.

## 2. Result

To begin, we assume that all the three-dimensional coordinates of localized fluorophores labeling the proteins of interest were previously determined, using Gaussian fitting or other methods. Numerous prior works and reviews cover details of the localization process, and we will not address it here[10,26]. Synapses are structurally unique, with presynaptic active zone (AZ) and postsynaptic density (PSD) always symmetrically aligned across the synaptic cleft, as is clearly visible under the electron microscope[27]. Therefore, when one AZ protein and one PSD protein are separately labeled, synapses can be efficiently identified as sandwich-shaped structures in the scatter plot of localizations[13]. The boundary of all the localizations of a protein in one synapse (which we term the “synaptic cluster”) can be defined with the DB-SCAN method[28].

### 2.1 The detection of high-density nanoclusters

#### Basic strategy

Within the synaptic cluster are frequently found further smaller clusters of protein, which we term “nanoclusters”[21] or nanodomains[19,20]. To automatically identify nanoclusters, we segmented the localizations within a cluster based on their local density, thus defining nanoclusters as groupings of particularly high-density localizations. Thus, accurately calculating the density threshold of nanoclusters is the most critical step. This is made more difficult because although the border of the synaptic cluster is often abrupt and steep, most synaptic proteins are not distributed with a high density contrast between nanoclusters and the background within the synaptic cluster, and the finite imaging resolution blurs apparent nanocluster borders. We took a strategy similar to the DB-SCAN method[28] to calculate the local density (*LD*) of localizations by counting the number of localizations within a certain distance (*d*) from each localization. To account for the variation in localization density across different synaptic clusters, we defined *d* as *T* □ *MNND* instead of a fixed value[19,21], where *MNND* is the <u>m</u>ean <u>n</u>earest <u>n</u>eighboring <u>d</u>istance of all localizations within the synaptic cluster, and *T* is a scalar multiplying factor. The appropriate value for *T* was determined empirically, as shown below. The threshold of local density for nanocluster detection was defined as *Mean*(*LD0*) + 4 x *Std*(*LD0*), where *LD0* is the local density of a randomized cluster with the same overall density as the synaptic cluster. This threshold represents the 99.95% confidence that the measured density differs from chance, and all localizations with a local density larger than this threshold were considered within nanoclusters. These localizations were then divided into sub-clusters with a “top-down” divisive strategy with a minimal peak-to-peak distance of 80 nm which is roughly the average size of synaptic nanoclusters[21]. Finally, to be counted as nanocluster, those sub-clusters had to meet a set of criteria, including number of localizations ≥8 localizations, which was derived empirically based on tests on our measured and simulated synapses to reduce the false-positives arising from repeated localizations of the same molecule.

#### Practical example

We labeled the postsynaptic scaffold protein PSD-95 in cultured hippocampal neurons with antibodies conjugated with Alexa647 and used 3D-STORM to map its distribution[10,29]. We integrated all the above processes into one MATLAB function *nanocluster_detection_3d.m* and used it to detect nanoclusters by varying the parameter *T* (Figure 1A). At the same time, we also randomized the distribution of localizations within the measured borders of the cluster and used the same function to detect any “false positive” nanoclusters (figure 1B). When *T* was small, i.e. local density was calculated within a smaller radius, the algorithm was too sensitive to the localization distribution within a very close vicinity and therefore the nanocluster number and their peak positions showed a larger variation; when *T* was too large, local densities were more washed out due to a larger averaging radius and therefore some nanoclusters were missing from the result, and the detected peak of a nanocluster (red dots in Figure 1A) did not represent the intuitive peak (Figure 1A, *T* = 8). At almost all values of *T*, the rate of detecting false-positives was very low (Figure 1C-D). For the example PSD-95 cluster, the nanocluster number was constant and the result matched our visual expectation with *T* 1∼3 (figure 1C). When results from more PSD-95 clusters (n = 59) were pooled together, the nanocluster number was stable for *T* 1∼2 and then started reducing with higher *T* (figure 1D). Based on these, we chose *T* of 2.5 for all our analysis.

**Figure 1.**
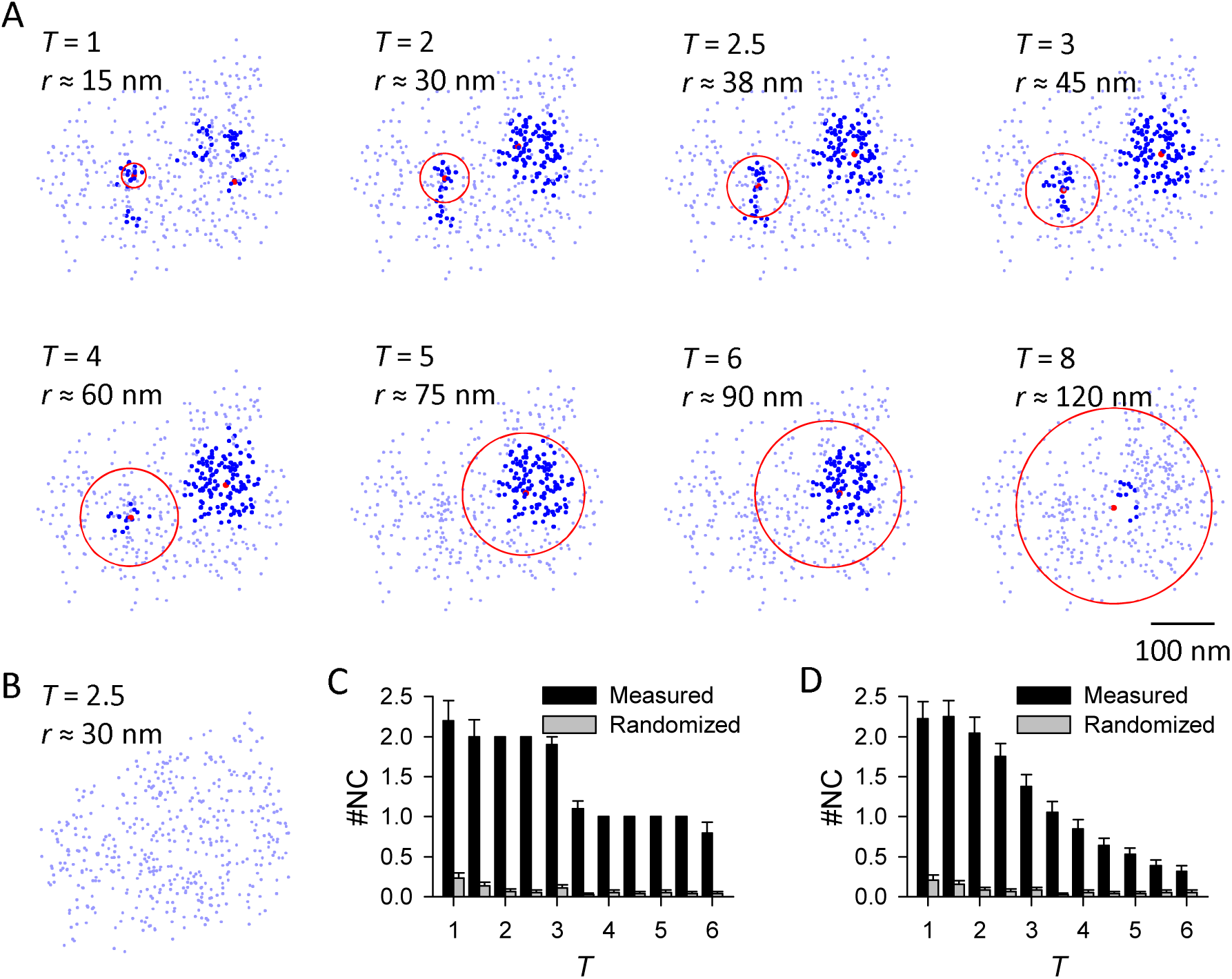
The detection of nanocluster and its robustness. **A.** Detected nanoclusters from the same PSD-95 cluster with different *Tra* values as labeled. PSD-95 localizations are shown in the enface angle, with the thick colored denoting localizations within nanoclusters and the red representing the peak of a nanocluster, i.e. localization with highest local density within a nanocluster. Note that the peak of a nanocluster is not necessarily around its center. Red circles represent the region within which the local density was calculated for nanocluster detection, with the radius (*r*)roughly calculated and labeled. Scale 100 nm. **B.** Typical example of homogenized cluster of the same one in **a**, with no nanocluster detected. **C.** Pooled results of detected nanocluster number in the example cluster in **a** from 20 rounds of computation. **D.** Pooled results of detected nanocluster number from 59 PSD-95 clusters. Note that in the full parameter space, the nanocluster number in measured protein was significantly higher than that of the randomized protein (p < 0.001 at all bins, one-way ANOVA on ranks with pairwise comparison procedures).

#### Discussion

The key parameter *T* was decided 2.5 as a tradeoff between two errors for this binary classification. First, we would like to reduce the false negative error, therefore we prefer a larger nanocluster number, which is favored by a lower *T* and a lower initial density threshold. Second, we want to minimize the false positive error, which requires a parameter set in the opposite direction. In our case, especially for the following protein enrichment analysis, the false positives would greatly affect the result by diluting the potential enrichment, while the impact of false negatives on enrichment result is minimal and the main risk is reducing the number of observations. Accordingly, we set the parameters to favor a lower false negative error, and as a result, we may have under-estimated the nanocluster numbers within synapses. In cases that the false negative is more critical, a lower *T* and a lower initial density threshold should be considered.

This method detects high local densities regardless of whether they arise from non-uniformed protein distribution or repetitive blinking of a single or a few fluorophores. The over-counting problem should be minimized during the sample preparation, imaging, and localization detection[30,31]. This algorithm can also obtain some detailed information of the nanoclusters including volume and the internal localization density. Please note that the nanocluster is not a discrete structure but a density gradient, and therefore this threshold-based algorithm would have a certain degree of arbitrariness. However, by applying the same set of detection parameters to different proteins or treatments, the method is sensitive enough to pick up differences in nanocluster number, volume or inside localization density[21].

For subsequent analysis below, it is not criticial what method is used to delineate the borders of subsynaptic nanoclusters. Indeed, there have been several methods well established for nanocluster detection in 2D data, including those based on DB-SCAN [19,20] or Voronoi tessellation [32], and these could be expanded to operate on 3D localizations. Due to the small synaptic cluster size and the relatively small number of localizations, the boundary effect around the edge of synaptic clusters would be a major challenge. However, the tessellation method has shown great potential in dealing with similar analysis[33].

### 2.2 Analysis of trans-synaptic protein alignment

To quantify the alignment of protein distributions across the synapse, we provide two independent methods: 1) 3D paired cross-correlation function (PCF) analysis to quantify the overall correlation of protein densities, and 2) a protein enrichment analysis to calculate the local protein density at positions opposing a given nanocluster on the other side of synapse[21,30]. Both methods were based on the assumption that the high densities within two proteins are distributed at similar positions within their own cluster, i.e. if the two synaptic clusters are overlapping, the two sets of high densities should be colocalized. However, the pre- and postsynaptic proteins are distributed at different sides of synaptic cleft with a distance of 50-200 nm[13]. Therefore, before performing these analyses of alignment, we first have to translate one cluster to overlap with the other without bias towards local densities.

#### 2.2.1 Overlapping pre- and postsynaptic clusters without bias towards local densities

##### Basic strategy

Though in EM images we cannot distinguish the protein identity or local density of specific proteins within active zones and postsynaptic densities, AZs and PSDs were always aligned well across the cleft[34], i.e., the AZ and PSD are two disc-shaped structures of the same size, paralleled with each other. Thus for proteins distributed within these two regions, such as RIM1/2 in AZ and PSD-95 in PSD, the space they take under 3D STORM should be similar in volume and shape[13,21]. Therefore, we could translate one cluster along a certain direction across a certain distance and get a good overlap with the other. Please note that this overlapping is for the general shape of the two synaptic clusters, as if aligning the edges of AZ and PSD in EM images. The effect of local high densities within the synaptic border should be minimized to avoid the circular argument logic fallacy in the following alignment analysis. Therefore, we set a density ceiling of ρ/4 for the density matrix of both clusters, where ρ is the average localization density within synaptic clusters. The magnitude and direction of the translation were then defined as the vector from the center to the peak in the cross-correlation space calculated with the density matrix of the two clusters[21].

##### Practical example

RIM1/2 and PSD-95 are key proteins in AZ and PSD[18, 47, 48], so we labeled these two proteins with antibodies conjugated with Alexa647 and Cy3, respectively, and used STORM to map their 3D distribution (Figure 2A)[9,10,29]. 3D density matrices were built for each cluster with a voxel size of 5 nm, and a 3D convolution with kernel size of 11 was applied to smooth the matrices (Figure 2B). As mentioned above, the maximal density in a matrix was set to a quarter of the average density to eliminate the major heterogeneity inside the synapse (Figure 2C). With these largely homogenized matrices, the vector could be calculated, and the two matrices could be translated to have an optimal overlap based on their general shapes (Figure 2D). Finally, the overlapping original protein densities were restored to the state of measurement for the following quantitative analysis (Figure 2E-F). This part of the process was coded into the same Matlab script with the following paired cross-correlation function.

**Figure 2.**
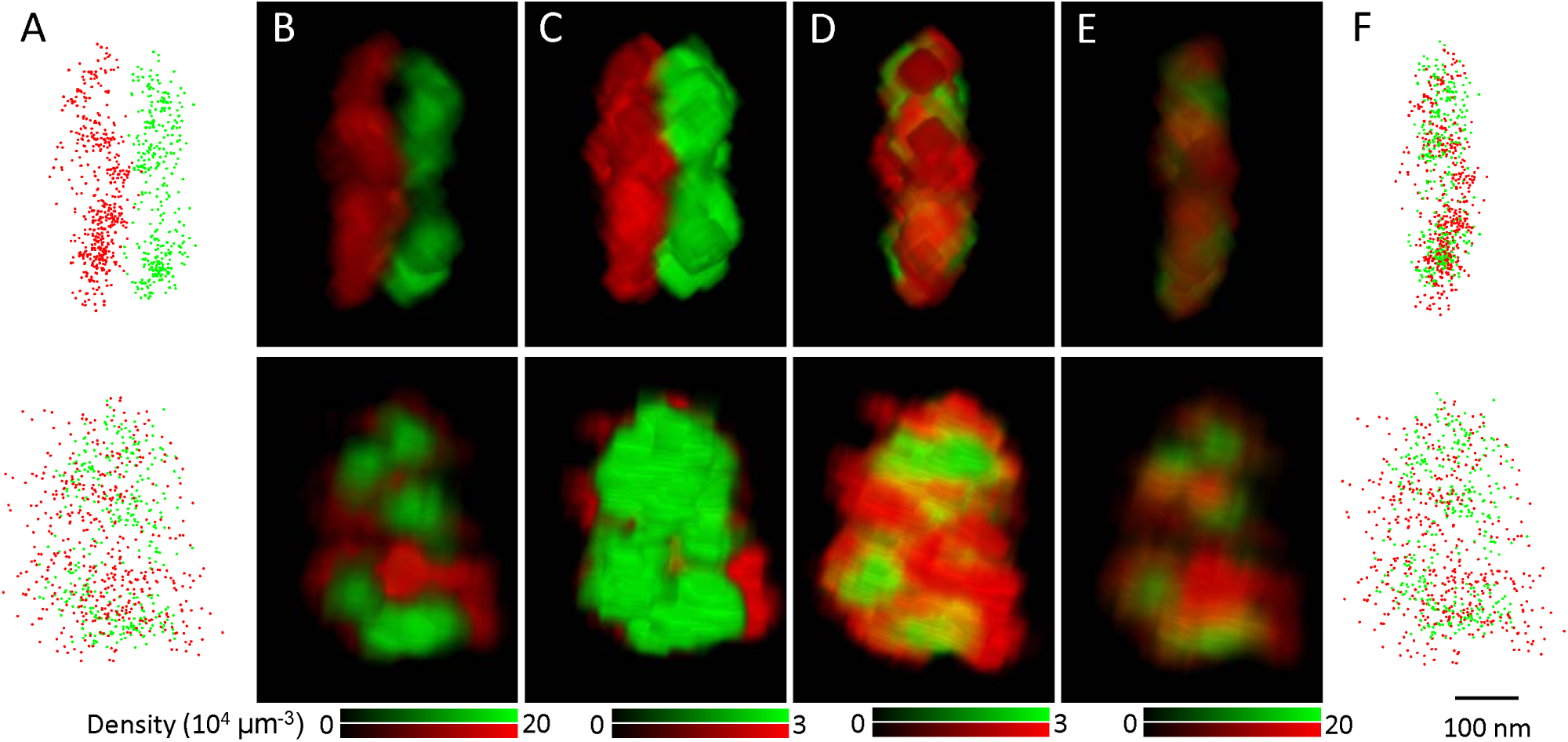
Translation of synaptic clusters to overlap pre- and postsynaptic clusters without bias towards local densities. **A.** Scatter plot of RIM1/2 (red) and PSD-95 (green) localizations with the side (top) and enface (bottom) view angles. **B-E.** Volume views of the original synaptic density matrix (**B**), matrix with a low density ceiling (**C**), matrix with density ceiling after the translation (**D**), and matrix with the original density after the translation (**E**). The density matrix was constructed with a voxel size of 5 nm and a 13×13 convolution was applied. Images were made with the 3D viewer plugin in Fiji ImageJ. **F.** Scatter plot of the two clusters fter the translation. Scale 100 nm.

##### Discussion

The translation was based on the assumption that the two clusters were proteins marking the AZ or PSD, that is, the protein structures attached to the synaptic membrane and thus fairly planar. So, extra caution should be taken for synaptic proteins with substantial presence away from the membrane, such as the protein synapsin which is associated with synaptic vesicles and thus fills almost the whole bouton[35,36].

In the defined function, we set one parameter to constrain the distance range of the translation (*distance* in *get_crosscorr_3d.m*). For major synaptic protein pairs, the range could be estimated based on previous STORM study by Dani et al[13]. Depending on the imaging system, there may be a channel registration error. Therefore, the range of translation distance should be expanded accordingly. Note that this error would be largely removed after the translation.

#### 2.2.2 Paired cross-correlation analysis

##### Basic strategy

Pair correlation function has been used to quantify heterogeneity within an organization or colocalization between systems[30,37,38]. The pair cross-correlation function, *g*(*r*), reports the increased probability of finding a similar localized signal in system 2 at a distance *r* away from a given localized signal in system 1.

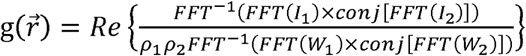

The function describes the cross-correlation of the two constructed density matrices (*I*_*1*_, *I*_*2*_) of the two sets of STORM localizations normalized with the cross-correlation of the two window functions (*W*_*1*_, *W*_*2*_) for *I*_*1*_ and *I*_*2*_, respectively. *W* has the value of 1 inside the corresponding cluster and 0 outside. The cross-correlation is tabulated in Matlab using Fast Fourier Transforms (FFTs), conj[] indicates a complex conjugate, ρ_*1*_ and ρ_*2*_ are the average densities of matrix *I*_*1*_ and *I*_*2*_ respectively, and Re indicates the real part. This normalization is critical as it removes all the effects coming from complex boundary shapes and makes the function account only for the internal density distributions within the two matrices. 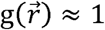 represents a random correlation between the two structures, and the colocalization of any high-density structures would result in 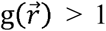. Since 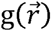 is theoretically symmetric to rotations, it can be averaged over all angles to obtain a one-dimensional g(*r*).

##### Practical example

We used the same RIM1/2 and PSD-95 cluster pair as example. Density matrix (*I*_*1*_ and *I*_*2*_) were built with a voxel size of 5 nm and the window functions *W*_*1*_ and *W*_*2*_ were defined as the same set of matrices with voxels set to 1 inside the cluster defined with an alpha shape (α =150 nm). In the three-dimensional 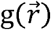 matrix (Figure 3A), the voxels in the center region showed significant higher values, which was clearer when it was circularly averaged to a one-dimensional distribution (Figure 3B). The g(*r*)between the measured proteins was significantly larger than 1 within a certain radius range (with ANOVA), suggesting the internal densities of RIM1/2 and PAD-95 at this synapse had a significant alignment.

**Figure 3.**
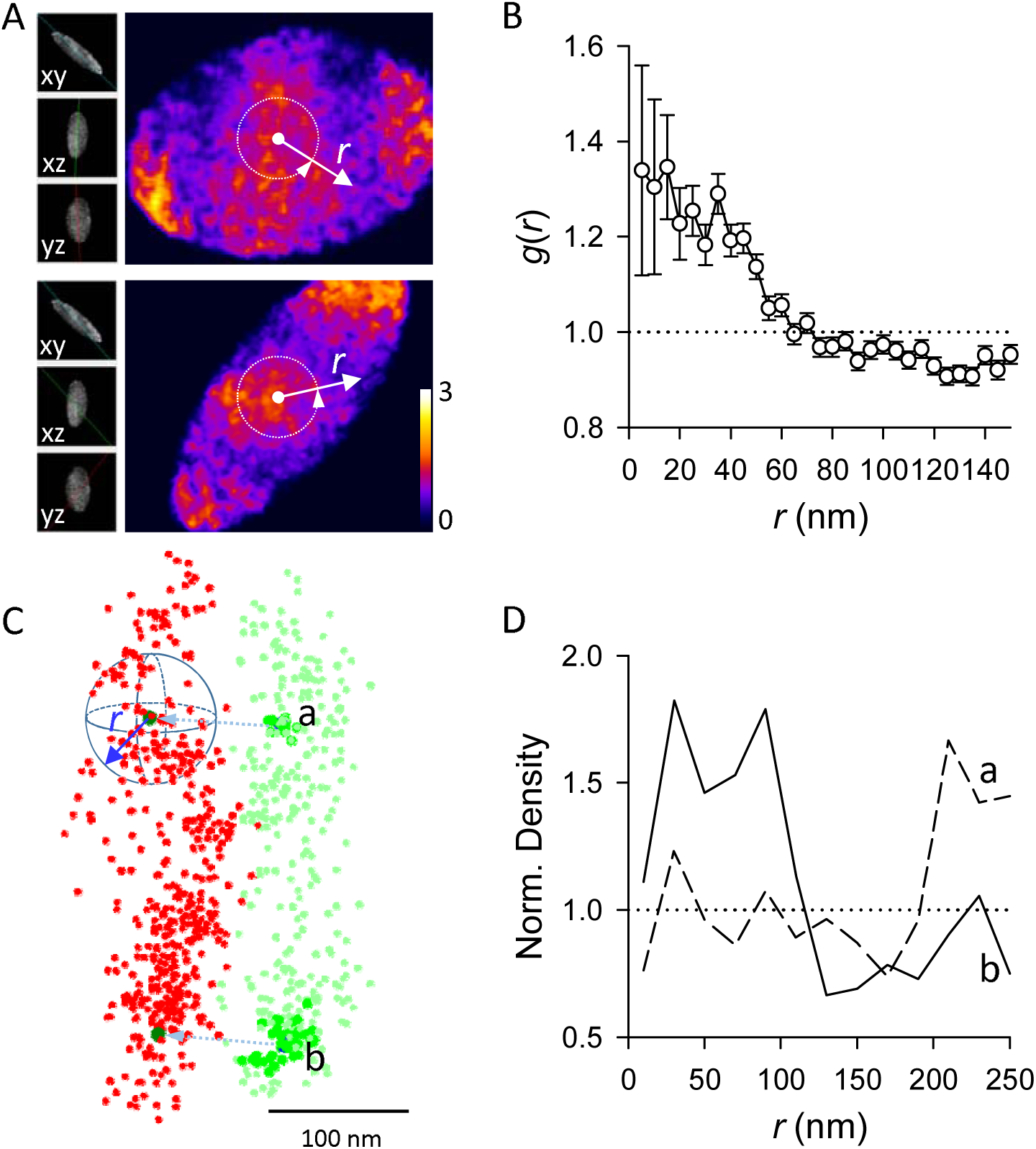
Paired cross-correlation and protein enrichment analysis. **A**. Two sections of the 3D paired cross-correlation matrix 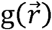. Inserts on the left represent the sectioning angles, and the color coded the normalized coefficients of paired correlation between density matrixes of RIM1/2 and PSD-95 after translation. Note the heated color near the center of matrix. Images were made with the volume viewer plugin in Fiji ImageJ. The color-coded are divided according to the degree of heat, and the left side shows different three-dimensional angles. r represents the size of the region between the various angles deviating from the best correlation value. **B.** The paired correlation function distribution g(*r*)averaged over all angles along distances from the center of matrix 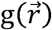, as shown with the white arrows in **A**. Note g(*r*)is significantly higher than 1 in within certain radius. **C.** Strategy of the protein enrichment analysis. From the projected peaks of PSD-95 nanoclusters (dark green), local density of RIM1/2 (red) was averaged over all angles along distances. In case of a positive alignment, a higher averaged density is expected at distances around 0. **D.** Spatial profile of normalized RIM1/2 density along distances from projected peaks of PSD-95 nanocluster **a** and **b**. Note the significant enrichment of RIM1/2 to nanocluster **b** but not to **a**.

All these computations were incorporated in a defined Matlab function *get_crosscorr_3d.m* together with the cluster translation in part 2.2.1. Besides the 3D coordinates of the two localization sets as inputs, the other parameters included voxel size, radius range to calculate g(*r*), cluster translation vector, and distance range of the translation. If the cluster translation vector was set as null ([] in Matlab), the function would calculate the vector as described in 2.2.1, which would be skipped if there was a valid input for cluster translation vector.

##### Discussion

The application of the paired cross-correlation method is not limited to the analysis of two neighboring or overlapped protein clusters, but can be expanded to many occasions for colocalization or alignment analysis such as the three-dimensional distribution of two proteins in a defined space with varied volumes. However, when distributions over a large volume are analyzed with a small voxel size, i.e. when there are a large voxel number in the constructed density matrix, running the function may be extremely memory-intensive in current versions of Matlab. In this case, an alternate computing strategy should be employed[39].

The paired cross-correlation method determines whether two three-dimensional clusters have correlated internal density structures[30,37]. It does not rely on the detection of high-density nanoclusters therefore won’t be affected by potential errors during nanocluster detection as discussed in part 2.1. For the same reason, this method cannot provide any detailed information about the alignment of individual nanoclusters and therefore it may not be sensitive enough to detect all potential alignment, especially when there are multiple high-density peaks. Ideally, we need an analysis that could test the alignment for each individual nanocluster – therefore, we developed the protein enrichment analysis.

#### 2.2.3 Protein enrichment analysis

##### Basic strategy

This analysis is based on the prediction that if the pre- and postsynaptic nanoclusters align across the cleft, the presence of a nanocluster on one side will predict a higher local protein density around its projected point on the other side. To quantitatively test this, we calculated the average local density of protein A over the distance from the projected peak of a protein B nanocluster. In case of a positive alignment, this curve would start from a local density significantly higher than the average at the small distance and then decay to the average.

##### Practical example

We explored the degree of RIM1/2 enrichment relative to the two PSD-95 nanoclusters in the same example synapse. The PSD-95 nanoclusters were detected as escribed in 2.1 and the PSD-95 cluster together with the two nanocluster peaks was projected to have a best overlap with RIM1/2 cluster as in 2.2.1. The number of RIM1/2 localizations were counted within binned distance ranges from the projected peaks (Figure 3C). To account for the impact of the cluster boundary on the density calculation, we randomized the RIM1/2 localizations within its cluster boundary and used the distribution of randomized localizations along the same distance to normalize the original distribution. If there was a RIM1/2 nanocluster aligning to a given PSD-95 nanocluster, we would expect a normalized density significantly larger than 1 within a certain small range of *r*, such as for nanocluster b in Figure 3C-D. This *r* range was determined as *r* < 60 nm based on pooled enrichment, and an enrichment index could be defined as the averaged density within the range for further statistical tests [21]. Otherwise, the normalized density would be around 1, which suggests a random distribution, as can be seen for nanocluster a in Figure 3C-D, or below 1, which would suggest that molecules are de-enriched in regions closely aligned with the nanocluster. To simplify the quantification, we defied an enrichment index by averaging the normalized density within 60 nm from the projected peak. The radius of 60 nm was chosen based on the fact that most key synaptic proteins are significantly enriched in the nanocolumn within this radius [21]. Whether RIM1/2 is enriched to a given PSD-95 nanocluster could be determined via comparison with the enrichment indices of multiple randomized RIM1/2 clusters. With this, the percentile of PSD-95 nanoclusters that were enriched with RIM1/2 could be quantified [21].

##### Discussion

Enrichment analysis and paired cross-correlation function are two independent tests on whether two clusters have spatially correlated internal density structures. While the paired cross-correlation compares the overall degree of correlation between internal distributions of two proteins, the enrichment analysis provides more detailed information by quantifying the local density of protein A relative to a defined B sub-region. While in our case of nanoscale alignment between high density peaks the sub-region of protein B is the defined nanoclusters, the same analysis could be easily adapted to other forms of sub-structures such as hollow spots or inverse density peaks depending on the demand of specific scientific questions. Since the enrichment analysis is based on the positions of sub-regions, the false-negatives of the nanocluster detection would have a great impact by diluting the distribution profile. Therefore, the analysis will benefit from stricter criteria on nanocluster detection.

Due to the discrete nature of SMLM data, the boundary effect could dominate the result in special occasions, especially when a ratiometric measurement is made. In our case, depending on the cluster shape and the position of the nanocluster, the valid volumes for some bins may be very small. Even though we randomized the cluster with a density 10 times the original, which was equivalent to averaging across 10 simulations, the numbers of randomized localizations within these volumes were still not representative, which would result in an extremely large ratio or even an invalid calculation. If a bin showed an infinite ratio, its neighboring bins usually suffered from this boundary effect. We exclude these bins when pooling the data to reduce potential contamination.

### 2.3 Automatic enface projection and averaging of synapses

While the enrichment analysis provides detailed spatial distribution of one protein along distances from defined points such as peaks of nanoclusters of the reference protein, it would be helpful if similar information can be represented as images. Here we present an automatic method to make a projection of the three-dimensional synaptic structure to a defined plane such as synaptic cleft to generate an enface view of the protein distribution. This projection would make it possible to average the enface profile of protein densities or even analyze the relative spatial distribution patterns of pre- and postsynaptic nanoclusters.

#### Basic strategy

With one presynaptic AZ protein and one postsynaptic PSD protein labeled with two fluorophores, a typical synapse would be a sandwich shape, or a flat disc after one set of localizations were translated to overlay moved and overlaid with the other (Figure 2). A plane parallel to the cleft can be defined by fitting all overlaid localizations (least square of the normal distance to the plane). The two-dimensional enface projection can be achieved with calculation of the projected coordinates of all localizations along the fitted plane.

#### Practical example

We use the same synapse as example. After overlaying the two cluster together as in Figure 2, we calculated the enface plane (Figure 4A) with the defined Matlab function *get_2D_projection.m* based on the *affine_fit.m* function by Adrien Leygue. The same function also yielded the projected 2D coordinates so we could generate the enface density map. To rule out the effect of cluster thickness along the projection direction on the local density, we randomized all localizations within the original clusters and performed a similar projection to get a map coding the thickness information. Using this as a normalizing factor, we obtained a map coding the related local density (Figure 4B).

**Figure 4.**
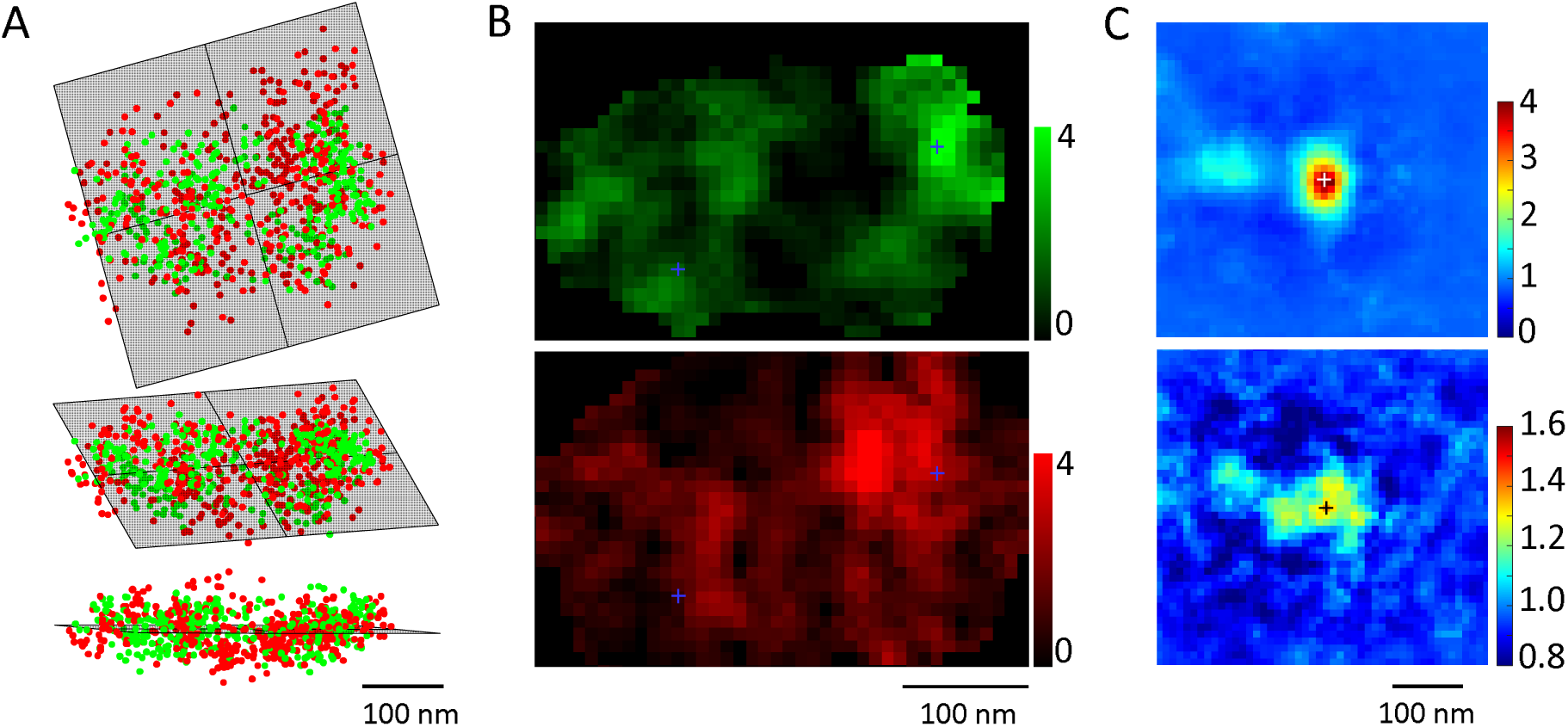
Automatic enface projection and averaging of synapses. **A**. Calculation of the enface plane by fitting all overlapped localizations after translation. RIM1/2 is in red, PSD-95 in green. Top shows the top-view, lower the side-view and middle the elevated view. **B**. Local density distribution after the projection. Crosses present the peaks of two PSD-95 nanoclusters. **C**. Averaged PSD-95 nanocluster in projection plane and the corresponding density distribution of RIM1/2 across 103 nanoclusters from 59 synapses. PSD-95 distribution was rotated to get the best fit of the internal distribution to the original template. Note the second nanocluster ∼120 nm from the center of the averaged nanocluster. RIM1/2 was rotated with the same angle as PSD-95 from the same synapse. Note the significant higher density around the averaging center.

To visualize the enface distribution of both RIM1/2 and PSD-95 around PSD-95 nanoclusters, we averaged both normalized density maps centered around the projected peaks of PSD-95 nanoclusters (crosses in Figure 4B-C). To further test whether there are secondary density structures around the nanoclusters, we performed a free 360-degree rotation around the nanocluster peak to find the best correlation with a template (density map around the first nanocluster for the first correlation, and the averaged density map for the following). Note that this correlation was computed in a similar way as in part 2.1.2 to eliminate the effect of cluster boundaries border, i.e. only the internal density structures mattered for the correlation. Meanwhile, to avoid any artifact created by the bordering effect, all values outside the synaptic cluster were replaced with 1 before averaging was performed. This process was incorporated in a defined function *get_bestfit_rotate.m*. While PSD-95 maps were freely rotated, the rotation angles of RIM1/2 maps were kept the same as that of PSD-95 in the same synapse to maintain the relative positioning of the two clusters. In the averaged density maps in Figure 4C, besides the center PSD-95 nanocluster, there was a secondary but weaker nanocluster, which is a result of freely-rotating averaging across the maps of synapses with 2 or more PSD-95 nanoclusters. The distance of ∼120 nm from the image center suggests an average distance between two neighboring nanoclusters for PSD-95. As expected, the averaged map of RIM1/2 also showed a significant higher density around the center, representing the enrichment between the two proteins.

#### Discussion

Theoretically, the translation vector should be vertical to the fitted plane and therefore we could simply use that vector to make projections. However, the vector was often contaminated by the two-channel registration error, so its direction was not as reliable for projection. The current method benefited from the fact that the registration error was largely reduced by the overlapping translation, and therefore provided not only a more accurate projecting direction but also less error for the 2D enrichment distribution, as demonstrated by the secondary PSD-95 nanocluster and the significant enrichment of RIM1/2 around the image center (Figure 4C).

Similar with the overlapping translation, the fitting of the enface plane assumed that the clusters were representing the disc shape of AZ or PSD. If one protein had a strong distribution outside these two specialized compartments, such as presynaptic synapsin and postsynaptic actin or mGluRs[36,40], it should not be included in fitting process. In this case, the enface plane could be fitted with only the other protein. Moreover, for large synapses, this algorithm may fail if complex border structure in the 2D projection of either presynaptic or postsynaptic shape dominates the projection.

## 3. Conclusions

Imaging with single molecule localization microscopy provides a wealth of information on subcellular structures and protein organizations which underly their specialized functions. To exploit them requires more detailed sophisticated quantitative analyses rather than image processing that most general software packages provide. We have described a set of detailed analytical methods on quantification of trans-synaptic alignment on three-dimensional STORM data and have made all the Matlab scripts available. Some could be easily adapted to analysis of other biological structures. We hope our methods could be helpful and inspiring for others to design automated and quantitative analysis on their SMLM data.

## Supporting information

Supplementary Matlab codes and example data. Zipped in rar format.

## ACKNOWLEDGMENTS

The authors acknowledge grant supports to AT (NARSAD Young Investigator Award, National Natural Science Foundation of China 31872759, and the USTC Youth Innovation Fund) and TAB (National Institutes of Health R01 MH080046 and NS090644). We thank Poorna Dharmasri for his comments on the manuscript.

## Author contributions

J.-H. C. and A.-H. T. prepared figures and drafted the manuscript; J.-H. C., T.A.B. and A.-H. T. finalized the figures and manuscript.

## Appendix A

Matlab scripts for all the analysis described.

## References

[1] J.E. Lisman, S. Raghavachari, R.W. Tsien, The sequence of events that underlie quantal transmission at central glutamatergic synapses, Nat Rev Neurosci. 8 (2007) 597–609. doi:10.1038/nrn2191.

[2] J. Kleinle, K. Vogt, H.R. Lüscher, L. Müller, W. Senn, K. Wyler, J. Streit, Transmitter concentration profiles in the synaptic cleft: an analytical model of release and diffusion, Biophysical Journal. 71 (1996) 2413–2426. doi:10.1016/S0006-3495(96)79435-3.

[3] S. Raghavachari, J.E. Lisman, Properties of Quantal Transmission at CA1 Synapses, Journal of Neurophysiology. 92 (2004) 2456–2467. doi:10.1152/jn.00258.2004.

[4] J. Lisman, S. Raghavachari, A Unified Model of the Presynaptic and Postsynaptic Changes During LTP at CA1 Synapses, Sci. STKE. 2006 (2006) re11–re11. doi:10.1126/stke.3562006re11.

[5] J. Lisman, Glutamatergic synapses are structurally and biochemically complex because of multiple plasticity processes: long-term potentiation, long-term depression, short-term potentiation and scaling, Phil. Trans. R. Soc. B. 372 (2017) 20160260. doi:10.1098/rstb.2016.0260.

[6] F.E. Bloom, G.K. Aghajanian, Cytochemistry of Synapses: Selective Staining for Electron Microscopy, Science. 154 (1966) 1575–1577. doi:10.1126/science.154.3756.1575.

[7] S.T. Hess, T.P.K. Girirajan, M.D. Mason, Ultra-High Resolution Imaging by Fluorescence Photoactivation Localization Microscopy, Biophysical Journal. 91 (2006) 4258–4272. doi:10.1529/biophysj.106.091116.

[8] E. Betzig, G.H. Patterson, R. Sougrat, O.W. Lindwasser, S. Olenych, J.S. Bonifacino, M.W. Davidson, J. Lippincott-Schwartz, H.F. Hess, Imaging Intracellular Fluorescent Proteins at Nanometer Resolution, Science. 313 (2006) 1642–1645. doi:10.1126/science.1127344.

[9] M.J. Rust, M. Bates, X. Zhuang, Sub-diffraction-limit imaging by stochastic optical reconstruction microscopy (STORM), Nature Methods. 3 (2006) 793–796. doi:10.1038/nmeth929.

[10] B. Huang, W. Wang, M. Bates, X. Zhuang, Three-Dimensional Super-Resolution Imaging by Stochastic Optical Reconstruction Microscopy, Science. 319 (2008) 810–813. doi:10.1126/science.1153529.

[11] H. Shroff, C.G. Galbraith, J.A. Galbraith, H. White, J. Gillette, S. Olenych, M.W. Davidson, E. Betzig, Dual-color superresolution imaging of genetically expressed probes within individual adhesion complexes, PNAS. 104 (2007) 20308–20313. doi:10.1073/pnas.0710517105.

[12] R. Jungmann, M.S. Avendaño, J.B. Woehrstein, M. Dai, W.M. Shih, P. Yin, Multiplexed 3D cellular super-resolution imaging with DNA-PAINT and Exchange-PAINT, Nat Meth. 11 (2014) 313–318. doi:10.1038/nmeth.2835.

[13] A. Dani, B. Huang, J. Bergan, C. Dulac, X. Zhuang, Superresolution Imaging of Chemical Synapses in the Brain, Neuron. 68 (2010) 843–856. doi:10.1016/j.neuron.2010.11.021.

[14] B. Dudok, L. Barna, M. Ledri, S.I. Szabo, E. Szabadits, B. Pinter, S.G. Woodhams, C.M. Henstridge, G.Y. Balla, R. Nyilas, C. Varga, S.-H. Lee, M. Matolcsi, J. Cervenak, I. Kacskovics, M. Watanabe, C. Sagheddu, M. Melis, M. Pistis, I. Soltesz, I. Katona, Cell-specific STORM super-resolution imaging reveals nanoscale organization of cannabinoid signaling, Nat Neurosci. 18 (2015) 75–86.

[15] M. Igarashi, M. Nozumi, L.-G. Wu, F.C. Zanacchi, I. Katona, L. Barna, P. Xu, M. Zhang, F. Xue, E. Boyden, New observations in neuroscience using superresolution microscopy, J. Neurosci. 38 (2018) 9459–9467. doi:10.1523/JNEUROSCI.1678-18.2018.

[16] K. Xu, G. Zhong, X. Zhuang, Actin, Spectrin, and Associated Proteins Form a Periodic Cytoskeletal Structure in Axons, Science. 339 (2013) 452–456. doi:10.1126/science.1232251.

[17] F. Pennacchietti, S. Vascon, T. Nieus, C. Rosillo, S. Das, S. Tyagarajan, A. Diaspro, A. del Bue, E.M. Petrini, A. Barberis, F.C. Zanacchi, Nanoscale molecular reorganization of the inhibitory postsynaptic density is a determinant of GABAergic synaptic potentiation, J. Neurosci. (2017) 0514–16. doi:10.1523/JNEUROSCI.0514-16.2016.

[18] T.J. Younts, H.R. Monday, B. Dudok, M.E. Klein, B.A. Jordan, I. Katona, P.E. Castillo, Presynaptic Protein Synthesis Is Required for Long-Term Plasticity of GABA Release, Neuron. 92 (2016) 479–492. doi:10.1016/j.neuron.2016.09.040.

[19] H.D. MacGillavry, Y. Song, S. Raghavachari, T.A. Blanpied, Nanoscale Scaffolding Domains within the Postsynaptic Density Concentrate Synaptic AMPA Receptors, Neuron. 78 (2013) 615–622. doi:10.1016/j.neuron.2013.03.009.

[20] D. Nair, E. Hosy, J.D. Petersen, A. Constals, G. Giannone, D. Choquet, J.-B. Sibarita, Super-Resolution Imaging Reveals That AMPA Receptors Inside Synapses Are Dynamically Organized in Nanodomains Regulated by PSD95, J. Neurosci. 33 (2013) 13204–13224. doi:10.1523/JNEUROSCI.2381-12.2013.

[21] A.-H. Tang, H. Chen, T.P. Li, S.R. Metzbower, H.D. MacGillavry, T.A. Blanpied, A trans-synaptic nanocolumn aligns neurotransmitter release to receptors, Nature. 536 (2016) 210–214. doi:10.1038/nature19058.

[22] H. Sakamoto, T. Ariyoshi, N. Kimpara, K. Sugao, I. Taiko, K. Takikawa, D. Asanuma, S. Namiki, K. Hirose, Synaptic weight set by Munc13-1 supramolecular assemblies,Nature Neuroscience. (2017) 1. doi:10.1038/s41593-017-0041-9.

[23] K.T. Haas, B. Compans, M. Letellier, T.M. Bartol, D. Grillo-Bosch, T.J. Sejnowski, M. Sainlos, D. Choquet, O. Thoumine, E. Hosy, Pre-post synaptic alignment through neuroligin-1 tunes synaptic transmission efficiency, ELife. (2018). doi:10.7554/eLife.31755.

[24] H. Chen, A.-H. Tang, T.A. Blanpied, Subsynaptic spatial organization as a regulator of synaptic strength and plasticity, Current Opinion in Neurobiology. 51 (2018) 147–153. doi:10.1016/j.conb.2018.05.004.

[25] L. Barna, B. Dudok, V. Miczán, A. Horváth, Z.I. László, I. Katona, Correlated confocal and super-resolution imaging by VividSTORM, Nat. Protocols. 11 (2016) 163–183. doi:10.1038/nprot.2016.002.

[26] S. Jia, J.C. Vaughan, X. Zhuang, Isotropic three-dimensional super-resolution imaging with a self-bending point spread function, Nat Photon. 8 (2014) 302–306. doi:10.1038/nphoton.2014.13.

[27] G.R. Phillips, J.K. Huang, Y. Wang, H. Tanaka, L. Shapiro, W. Zhang, W.-S. Shan, K. Arndt, M. Frank, R.E. Gordon, M.A. Gawinowicz, Y. Zhao, D.R. Colman, The Presynaptic Particle Web: Ultrastructure, Composition, Dissolution, and Reconstitution, Neuron. 32 (2001) 63–77. doi:10.1016/S0896-6273(01)00450-0.

[28] M. Ester, H.-P. Kriegel, J. Sander, X. Xu, A Density-Based Algorithm for Discovering Clusters in Large Spatial Databases with Noise, Proc. 2nd International Conference on Knowledge Discovery and Data Mining. (1996) 226–231.

[29] S. van de Linde, A. Löschberger, T. Klein, M. Heidbreder, S. Wolter, M. Heilemann, M. Sauer, Direct stochastic optical reconstruction microscopy with standard fluorescent probes, Nat. Protocols. 6 (2011) 991–1009. doi:10.1038/nprot.2011.336.

[30] S.L. Veatch, B.B. Machta, S.A. Shelby, E.N. Chiang, D.A. Holowka, B.A. Baird, Correlation Functions Quantify Super-Resolution Images and Estimate Apparent Clustering Due to Over-Counting, PLOS ONE. 7 (2012) e31457. doi:10.1371/journal.pone.0031457.

[31] N. Durisic, L.L. Cuervo, M. Lakadamyali, Quantitative super-resolution microscopy: pitfalls and strategies for image analysis, Current Opinion in Chemical Biology. 20 (2014) 22–28. doi:10.1016/j.cbpa.2014.04.005.

[32] F. Levet, E. Hosy, A. Kechkar, C. Butler, A. Beghin, D. Choquet, J.-B. Sibarita, SR-Tesseler: a method to segment and quantify localization-based super-resolution microscopy data, Nature Methods. 12 (2015) 1065–1071. doi:10.1038/nmeth.3579.

[33] L. Andronov, J. Michalon, K. Ouararhni, I. Orlov, A. Hamiche, J.-L. Vonesch, B.P. Klaholz, 3DClusterViSu: 3D clustering analysis of super-resolution microscopy data by 3D Voronoi tessellations, Bioinformatics. 34 (2018) 3004–3012. doi:10.1093/bioinformatics/bty200.

[34] T. Biederer, P.S. Kaeser, T.A. Blanpied, Transcellular Nanoalignment of Synaptic Function, Neuron. 96 (2017) 680–696. doi:10.1016/j.neuron.2017.10.006.

[35] O. Bloom, E. Evergren, N. Tomilin, O. Kjaerulff, P. Löw, L. Brodin, V.A. Pieribone, P. Greengard, O. Shupliakov, Colocalization of synapsin and actin during synaptic vesicle recycling, The Journal of Cell Biology. 161 (2003) 737–747. doi:10.1083/jcb.200212140.

[36] J.-H. Tao-Cheng, Activity-related redistribution of presynaptic proteins at the active zone, Neuroscience. 141 (2006) 1217–1224. doi:10.1016/j.neuroscience.2006.04.061.

[37] P. Sengupta, T. Jovanovic-Talisman, D. Skoko, M. Renz, S.L. Veatch, J. Lippincott-Schwartz, Probing protein heterogeneity in the plasma membrane using PALM and pair correlation analysis, Nat Meth. 8 (2011) 969–975. doi:10.1038/nmeth.1704.

[38] M.B. Stone, S.L. Veatch, Steady-state cross-correlations for live two-colour super-resolution localization data sets, Nature Communications. 6 (2015) 7347. doi:10.1038/ncomms8347.

[39] Y. Yin, W.T.C. Lee, E. Rothenberg, Ultrafast data mining of molecular assemblies in multiplexed high-density super-resolution images, Nature Communications. 10 (2019) 119. doi:10.1038/s41467-018-08048-2.

[40] Z. Nusser, E. Mulvihill, P. Streit, P. Somogyi, Subsynaptic segregation of metabotropic and ionotropic glutamate receptors as revealed by immunogold localization, Neuroscience. 61 (1994) 421–427. doi:10.1016/0306-4522(94)90421-9.

